# Population bottleneck has only marginal effect on fitness evolution and its repeatability in dioecious *C. elegans*

**DOI:** 10.1101/2021.10.07.463474

**Authors:** Karen Bisschop, Thomas Blankers, Janine Mariën, Meike T. Wortel, Martijn Egas, Astrid T. Groot, Marcel E. Visser, Jacintha Ellers

## Abstract

The predictability of evolution is expected to depend on the relative contribution of deterministic and stochastic processes. This ratio is modulated by effective population size. Smaller effective populations harbor less genetic diversity and stochastic processes are generally expected to play a larger role, leading to less repeatable evolutionary trajectories. Empirical insight into the relationship between effective population size and repeatability is limited and focused mostly on asexual organisms. Here, we tested whether fitness evolution was less repeatable after a population bottleneck in obligately outcrossing populations of *Caenorhabditis elegans*. Replicated populations founded by 500, 50, or 5 individuals (no/moderate/strong bottleneck) were exposed to a novel environment with a different bacterial prey. As a proxy for fitness, population size was measured after one week of growth before and after 15 weeks of evolution. Surprisingly, we found no significant differences among treatments in their fitness evolution. Even though the strong bottleneck reduced the relative contribution of selection to fitness variation, this did not translate to a significant reduction in the repeatability of fitness evolution. Thus, although a bottleneck reduced the contribution of deterministic processes, we conclude that the predictability of evolution may not universally depend on effective population size, especially in sexual organisms.

## INTRODUCTION

The predictability of organismal evolution lies at the heart of our understanding of evolutionary theory (Losos, 2017; Orr, 2005; Stern and Orgogozo, 2008) and is now rapidly gaining broader interest, as it provides the basis for emergent evolutionary forecasting applications (Blount et al., 2018; Lässig et al., 2017; Nosil et al., 2020; Wortel et al., 2021). A major question is whether evolution follows mostly deterministic trajectories, i.e. repeatable in time and space, or whether non-deterministic processes dominate. Studies involving replicated populations in nature, such as repeated evolution of ecotypes, repeated colonization of islands, or repeated host race formation, have provided support for both parallel and non-parallel evolutionary trajectories (Colosimo et al., 2005; Elmer et al., 2014; Losos and Ricklefs, 2009; Nosil et al., 2002). Similarly, replicated evolutionary experiments in the laboratory have shown that short-term (one or a few dozen of generations) and long-term evolution can be both repeatable and not repeatable (Barrick et al., 2020; Blount et al., 2018; Graves et al., 2017; Travisano et al., 1995). These variable outcomes likely result from different relative contributions from determinism and stochasticity: in the absence of chance events, evolution is highly repeatable (Lässig et al., 2017). It is therefore important to understand which properties of the natural world influence the balance between deterministic and stochastic processes and the extent to which this affects the repeatability of evolution.

In experimental evolution studies that test the repeatability of evolution, most adaptation is fueled by selection on standing genetic variation rather than mutation (Barrett and Schluter, 2008). Thus, the dominant deterministic force in experimental evolution is positive selection and the dominant stochastic force is genetic drift. A critical factor that determines the relative importance of selection versus drift is the effective population size. In the Wright-Fisher model, changes in allele frequencies are strongly determined by drift if the effective population size is small relative to the strength of selection (Crow and Kimura, 1970; Fisher, 1923; Wright, 1931). However, since several aspects of fitness evolution are affected by population size, theory is ambivalent about the expected relationships between population size and the effects of drift versus selection, and thus the repeatability of fitness evolution. For example, we would expect effective population size to be positively related to average fitness, as in larger populations the efficacy of selection is higher (Kimura, 1983) and the effects of drift are reduced (Kimura et al., 1963; Willi et al., 2006). However, in very large populations the multitude of genotypic combinations opens up additional evolutionary trajectories, which may decrease the repeatability relative to moderately large populations (Szendro et al., 2013). In small populations, evolutionary trajectories are more heterogeneous, because there is a smaller chance that the most beneficial variant will get fixed, so that fitness effects of substitutions are more variable (De Visser and Rozen, 2005; Rozen et al., 2008). More variable effects of substitutions can allow small populations to obtain higher fitness peaks than large populations, e.g. if the most beneficial variants are fixed by chance (Rozen et al., 2008; Whitlock et al., 1995).

Sex (recombination) further complicates (the repeatability of) fitness evolution, because sex can increase the likelihood of evolution towards higher fitness in both large and small populations, but high recombination rates may prevent adaptation, especially in large populations (Weissman et al., 2010). Therefore, especially for sexual populations, theoretical research indicates that the conditions under which larger populations display adaptive advantage and higher evolutionary predictability over smaller populations depend on detailed genetic knowledge of the organism under study.

In evolutionary experiments, effective population sizes are generally varied by different numbers of clonally reproducing cells or by bottlenecking ancestral populations of sexually reproducing organisms (Kawecki et al., 2012). A population bottleneck randomly selects a subset of the available genotypes and thus reduces genetic diversity; effective population sizes will remain low after a bottleneck for an extended period, because genetic diversity is lost much more quickly due to drift in small populations compared to increasing diversity due to new mutations (Kimura et al., 1963; Wright, 1931). Empirical data generally show that fitness in evolved small populations is more variable (less repeatable) and on average lower compared to large populations (Lachapelle et al., 2015; Rozen et al., 2008; Weber, 1990; Wein and Dagan, 2019; Windels et al., 2021), with some exceptions (Miller et al., 2011; van Dijk et al., 2017). However, in these studies population sizes were still in the thousands of breeding/clonally reproducing individuals or effective populations sizes cannot be disentangled from census population sizes. The limited exploration of sexual species and populations with strongly reduced effective sizes leaves an important gap in our knowledge about the role of population size in the adaptive potential and repeatability of fitness evolution.

Here we tested whether adaptation is faster and more repeatable in populations with large versus small effective sizes in an obligatory outbreeding line of the bacterivorous nematode *C. elegans*. We exposed replicate populations to a novel environment with a different bacterial prey and measured fitness as the population size after one week on the novel food source prior to and after 15 weeks of exposure to the novel conditions. To disentangle the impact of effective population size from the effects of census population size, we started the experiment with either 500 nematodes (at expected 1:1 sex ratio) from a large and genetically variable ancestral population or 500 nematodes derived from the same ancestral population subjected to a moderate or strong bottleneck. These experiments addressed two questions: i) What is the effect of a population bottleneck on the average and maximum fitness after selection?; ii) What is the effect of population bottlenecks on the repeatability of fitness evolution?

## METHODS

### Study species and creation of the bottlenecked populations

We performed experimental evolution using the *C. elegans* D00 population from the Teotónio lab (IBENS, Paris), which is a multiparent intercrossed population that is obligatorily outcrossing (Noble et al., 2017; Theologidis et al., 2014). Sex ratio in dioecious *Caenorhabditis* species and lines is expected to be 1:1 (Gray and Cutter, 2014). The D00 ancestral population was expanded on Nematode Growth Medium (NGM) (Stiernagle, 2006) plates seeded with *E. coli* OP50 at 20°C and divided in aliquots. Care was taken during this phase to maintain sufficiently large population sizes. To create the bottlenecked populations, five aliquots of the starting population were thawed and expanded at 20°C with *E. coli* OP50 as a food source. After six days, from each expanded population 5 or 50 female nematodes were transferred to a separate plate for the strong bottleneck and moderate bottleneck treatments, respectively. The females were chosen randomly from all available non-gravid females on a plate. We only selected females to avoid variation among bottleneck replicates by stochastically sampling different numbers of males and females. These bottlenecked populations were then grown for six days on NGM *E. coli* at 20°C before collecting the nematodes in Eppendorf tubes. Simultaneously with this expansion, for the “no bottleneck” treatment the other five aliquots from the ancestral population were thawed and expanded on NGM *E. coli* at 20°C for six days (Fig. S1). After expansion, nematodes were collected and for each replicate the population density was estimated to calculate the transfer volume required to transfer 500 worms. In this way, no-bottleneck and bottleneck treatments always started with 500 nematodes in an expected 1:1 sex ratio, but for the no-bottleneck treatment, these 500 nematodes were offspring of diverse ancestral populations, while for the bottleneck treatments, these 500 nematodes were offspring of 50 or only 5 founder females. Each treatment had five replicates, each consisting of three plates (to avoid the loss of a replicate if one plate would fail due to contamination or human error) that were each initiated with 500 nematodes (Fig. S1). All populations of *C. elegans* were maintained on plates (⌀ 9 cm) with ±12 mL NGM.

### Novel conditions for experimental evolution

During experimental evolution, *C. elegans* populations were grown on *Bacillus megaterium* (DSM No. 509). *Bacillus megaterium* is a poor food that results in impeded growth rates and, when given a choice, *C. elegans* avoids patches with *B. megaterium* (Shtonda and Avery, 2006).

In addition to the novel diet, the temperature was lowered to 16°C, which was done for experimental feasibility. The temperature reduction to 16°C may affect metabolic functions and defense pathways (Gómez-Orte et al., 2018) and therefore constituted an additional selection pressure. Moreover, 16S amplicon sequencing data from empty NGM plates revealed unexpected contamination of the plates (mainly bacteria from the genera *Serratia* and *Pseudomonas*). Even though empty plates did not reveal any visual bacterial growth at room temperature and the same plates were used for all the replicates in the different treatments, this contamination may have induced an unanticipated additional selection pressure. Since the effects of the three perturbations (novel food source, novel temperature, and plate contaminants) cannot be disentangled, they are considered together as the novel conditions.

### Experimental set-up

The experiment was initiated on fresh NGM plates with a lawn of *B. megaterium* and 500 nematodes per plate per replicate. Every week each plate was replaced by a new plate while 500 nematodes were transferred by washing the plates, mixing the three plates per replicate, estimating the density of nematodes and pipetting the necessary volume to the new plate. At the beginning (week 0) and end (week 15) of the experiment, large samples of the nematode populations were cryopreserved at −80°C until they were needed for fitness assessments.

### Fitness assessment

As a fitness proxy for each replicate nematode population, we estimated the population size achieved after one week of growth on *B. megaterium* at 16°C (Fig. S2) starting from 500 individuals, drawn randomly from the week 0 or 15 population. Frozen week 0 and week 15 populations were thawed simultaneously and 250 µL of each population was expanded on NGM *E. coli* at 20°C for one week to create a common garden. After expansion, 500 nematodes were transferred to each of three fitness assessment plates (NGM *B. megaterium* at 16°C) for each of the replicates. The size of the population after one week of growth (i.e. the fitness proxy) was extrapolated from counts in droplets of 5 µL (Fig. S2). These extrapolations were strongly correlated to counts obtained using a flow cytometer (Fig. S3).

All replicates were measured in triplicate. However, for some replicates the fitness was assessed on two or three separate fitness assessment days (leading to 6 or 9 measurements respectively, Table S1), which was possible as populations were frozen in several Eppendorf tubes at the same week. Additional details are available in the Supplementary Methods file.

### Data analysis

All statistical analyses were done in R version 4.1.0 with packages ‘emmeans’ v. 1.6.1 (Lenth, 2021), ‘MuMIn’ v. 1.43.17 (Bartoń, 2020), ‘lawstat’ v. 3.4 (Gastwirth et al., 2020), ‘lme4’ v. 1.1.27.1. (Bates et al., 2014) and plotting was done using ‘ggplot2’ v. 3.3.5 (Wickham, 2016).

#### Effect of population bottlenecks on average and maximum fitness across replicates

We tested for a significant increase in fitness (extrapolated population size after seven days of growth on *B. megaterium*) using linear mixed effect models. The dependent variable was the extrapolated population size and the fixed effects were week, treatment, and their interaction; replicate and fitness assessment day were included as random variables because the levels of these variables are random relative to the population they come from (typical for variables such as time/experimenter/measurement, etc.) and we care about their effects as a whole and not per level (Snijders and Bosker, 1999). Pairwise comparisons among fixed effect factor levels were done using Tukey’s method and we performed the Levene’s Test of Equality of Variances to investigate whether the variance of the fitness differed among treatments both before and after selection. We expected similar initial fitness for all treatments and lower final mean fitness and higher variance in fitness for the bottlenecked versus the no bottleneck populations.

We also investigated potential differences between treatments in the mean selection response (the difference between week 15 and week 0 per replicate). We used an Ordered Heterogeneity Test (Rice and Gainest, 1994), because we expected an order in the selection response: stronger response for the treatment without bottleneck compared to the bottlenecked populations and also a stronger response for the moderate bottleneck compared to the strong bottleneck. We also included alternative hypotheses where two treatments were equal, but different from the third treatment (Neuhäuser and Hothorn, 2006). The Ordered Heterogeneity Test was based on a Kruskal-Wallis rank sum test.

#### Effect of population bottlenecks on the repeatability of fitness evolution

Repeatability of fitness evolution was measured and compared between the no bottleneck treatment, the moderate bottleneck treatment, and the strong bottleneck treatment following two criteria: (i) the variance among realized selection responses across replicates within treatments (lower variance implies higher repeatability) and (ii) variance partitioning among effects from selection and chance (more relative variance attributed to selection implies higher repeatability).

i. We compared differences in the variance of the selection response across the five replicates among treatments using an Ordered Heterogeneity Test (Neuhäuser and Hothorn, 2006; Rice and Gainest, 1994) based on Levene’s Test of Equality of Variances. We expected that populations that underwent a (stronger) bottleneck had more variance in fitness across replicates.
ii. To test whether the relative contribution from chance and selection depended on the presence and strength of a bottleneck, we partitioned variance in effects of selection (variance between before experimental evolution and after 15 weeks of experimental evolution), chance (variance among replicates), and measurement error (variance among fitness assessment measurements nested within replicate) for each of the three treatments. We fitted a nested ANOVA model and extracted the mean squares to obtain relative proportions of the mean squares for each effect. The data were unbalanced due to variation in the number of fitness assessment days done per replicate. We therefore subsampled the data within treatments for all possible combinations of fitness assessment days. This resulted in 108 data sets for the “no bottleneck” treatment (3 × 3 × 3 × 2 × 2 combinations of fitness assessment days) and 8 data sets (2 × 2 × 2 × 1 × 1) for the “moderate” and “strong bottleneck” treatments. For each data set we calculated variance proportions, which can be compared across treatments because they come from balanced designs. We expected that populations that underwent a (stronger) bottleneck had more variance in fitness attributable to chance (drift) relative to selection.

## RESULTS

### Effect of population bottlenecks on average and maximum fitness across replicates

Before selection to the novel conditions, the populations under the moderate bottleneck treatment had a significantly lower fitness than the populations not exposed to a bottleneck (*t-* ratio = −4.779 and *p*-value < 0.0001) and the populations that underwent a strong bottleneck (*t*-ratio = −7.152 and *p-*value <0.0001, Table 1). Also, the variance in fitness before selection was significantly smaller in the moderate bottleneck treatment compared to the treatment without bottleneck (Levene’s test statistic = 89.676 and *p-*value < 0.0001) and the strong bottleneck treatment (Levene’s test statistic = 11.747 and *p*-value = 0.0019). After selection, fitness was higher in all treatments compared to the start of the experiment (Fig. 1A). However, the average fitness after 15 weeks of selection did not differ significantly among bottleneck treatments (Table 1). Similarly, no significant differences in variances in fitness across replicates were found among treatments (Ordered Heterogeneity test statistic = 0.584 and *p*-value = 0.5599). Both before and after evolution and across all treatments, there was variation among fitness assessment days (Fig. S4).

**Table 1:** Effect of population bottleneck after selection across replicates. We fitted a linear mixed effects model on the logarithm of the extrapolated counts. R^2^ = 0.80; predicted R^2^ = 0.75 A) Model summary statistics for fixed effects. B) Pairwise comparisons of the least square means within treatments and within weeks.

A) Summary of fixed effects
|  | Estimate | Std. Error | t value | Approximate P(>\|t\|) |
| --- | --- | --- | --- | --- |
| (Intercept, i.e. week 0, no bottleneck) | 7.37 | 0.30 | 24.97 | <0.0001 |
| week 15 | 2.22 | 0.22 | 10.23 | <0.0001 |
| moderate bottleneck | -1.33 | 0.43 | -3.10 | 0.0019 |
| strong bottleneck | 0.84 | 0.43 | 1.96 | 0.0499 |
| week 15 x moderate bottleneck | 0.94 | 0.37 | 2.57 | 0.0102 |
| week 15 x strong bottleneck | -0.94 | 0.37 | -2.57 | 0.0105 |

B) Pairwise comparisons *within treatments between weeks*
| Contrast | Estimate | SE | df | t-ratio | p-value |
| --- | --- | --- | --- | --- | --- |
| no bottleneck (week 0 - 15) | -2.22 | 0.24 | 123 | -9.23 | < 0.0001 |
| moderate bottleneck (week 0 - 15) | -3.16 | 0.31 | 123 | -10.07 | < 0.0001 |
| strong bottleneck (week 0 - 15) | -1.28 | 0.31 | 123 | -4.07 | 0.0205 |

*within weeks between treatments*
| Contrast | Estimate | SE | df | t-ratio | p-value |
| --- | --- | --- | --- | --- | --- |
| week 0 (no - moderate) | 1.33 | 0.45 | 123 | 2.98 | 0.0347 |
| week 0 (no - strong) | -0.84 | 0.45 | 123 | -1.88 | 0.1948 |
| week 0 (mod. - strong) | -2.18 | 0.36 | 123 | -5.97 | < 0.0001 |
| week 15 (no - moderate) | 0.39 | 0.38 | 123 | 1.04 | 0.5652 |
| week 15 (no - strong) | 0.10 | 0.38 | 123 | 0.26 | 0.9635 |
| week 15 (mod. strong) | -0.29 | 0.34 | 123 | -0.87 | 0.6687 |

**Figure 1:**
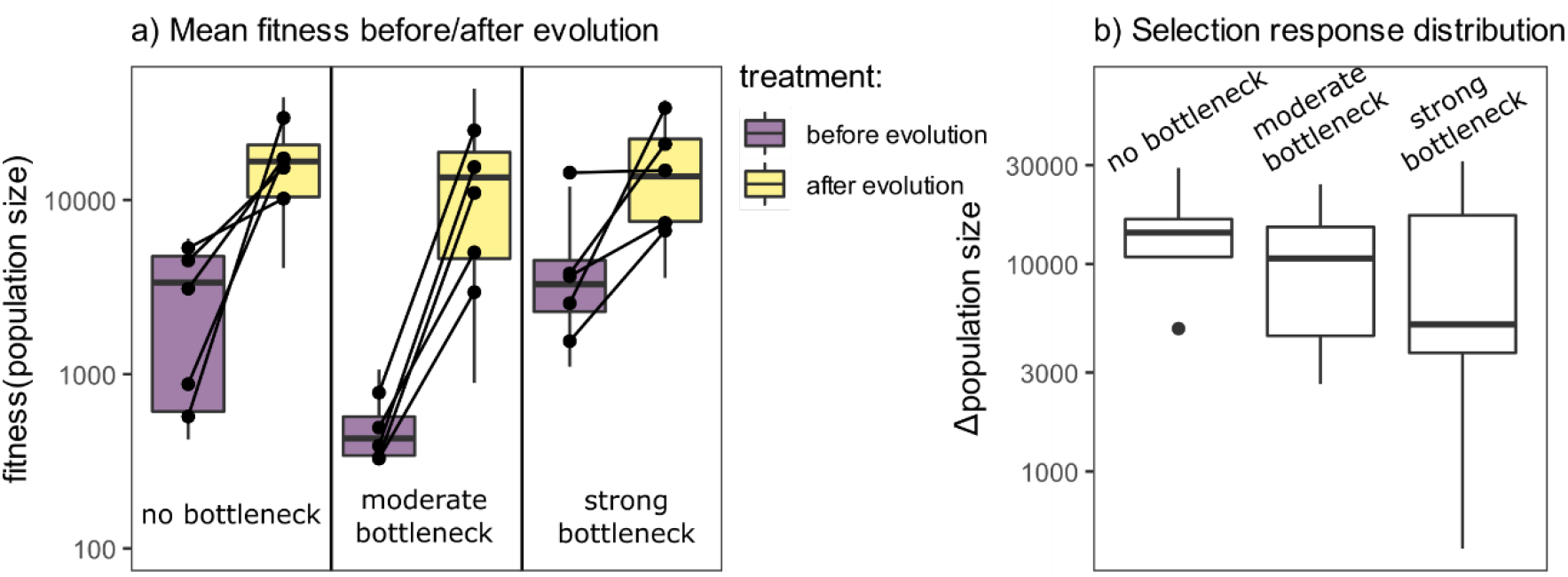
(a) Fitness (as population sizes measured in fitness assay after seven days of growth on B. megaterium) before and after selection. Box-and-whisker plots show distributions across measurements (three measurements per assessment day, per replicate, between 1 and 3 assessment days per replicate). Black lines connect replicate averages (across measurements) before and after selection. **(b) Selection response**, i.e. the difference in population size after 7 days of growth on *B. megaterium* between week 0 (unadapted) populations and week 15 (putatively adapted) populations.

The average selection response, measured as the difference in fitness between the week 0 and week 15 sample of a replicate, was highest in the no bottleneck treatment (mean = 15,097 and median = 14,184), but not significantly different from the selection response in the moderate bottleneck treatment (mean = 11,446 and median = 10,664) and the strong bottleneck treatment (mean = 11,541 and median = 5,118; Fig. 1B), based on the Ordered Heterogeneity Test (*r*_*s*_*P*_*c*_ statistic = 0.327, *P* = 0.228). The alternative hypothesis, that the bottleneck treatments did not differ from each other but had a lower selection response than the treatment without bottleneck, was also not significant (*r*_*s*_*P*_*c*_ statistic = 0.573, *P* = 0.073). Neither maximum fitness or maximum selection response across replicates were lower in the moderate bottleneck or strong bottleneck treatment compared to the no bottleneck treatment (Fig. 1).

### Effect of population bottlenecks on the repeatability of fitness evolution

The variance in the response to selection was not higher in the no bottleneck treatment compared to the moderate bottleneck treatment or the strong bottleneck treatment, based on the Ordered Heterogeneity Test (*r*_*s*_*P*_*c*_ statistic = 0.383, *P* = 0.182; Fig. 1B). The alternative hypothesis, with only a larger variance in the strong bottleneck compared to the other two treatments, was not supported either (*r*_*s*_*P*_*c*_ statistic = 0.671, *P* = 0.056).

When we compared the distributions of relative proportions of variance explained by error, chance, and selection between the no, moderate, and strong bottleneck treatments, we found that in the strong bottleneck treatment the relative proportion of the chance effect was always larger compared to the no bottleneck treatment and the moderate bottleneck treatment (Fig. 2). Because the variance attributable to measurement error was not different among treatments, the relative variance attributable to selection was thus lower after a strong bottleneck.

**Figure 2:**
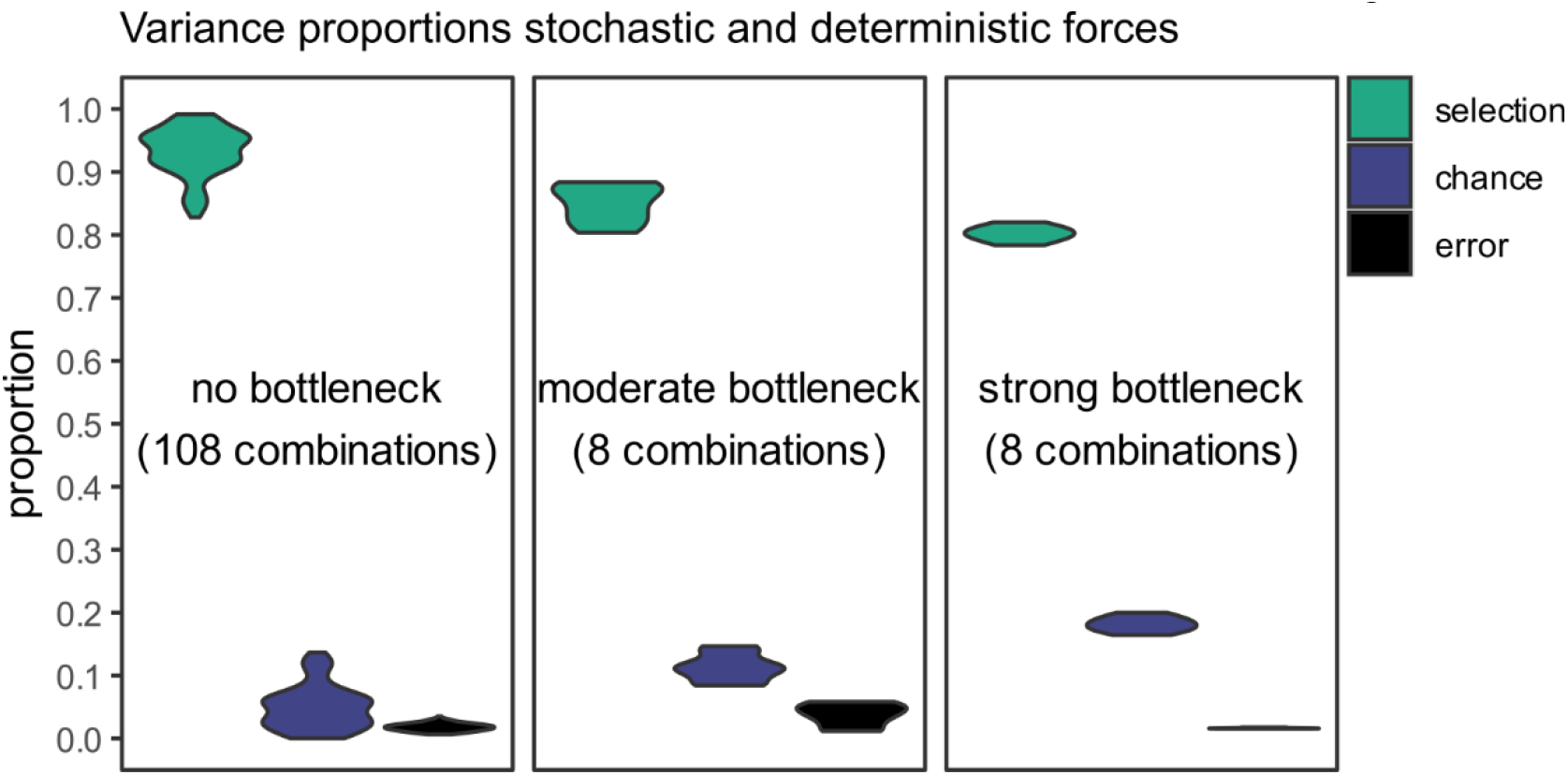
Variance partitioning between selection, chance, and error. The data was subset on the level of technical replicates (measurement occasions) to balance data prior to calculating mean squares and to explore variance proportions across all possible subsets. A higher number of measurements days for the replicates of the no bottleneck scenario results in more combinations of different measurement days compared to the moderate and strong bottleneck replicates. Violin plots indicate the spread of the proportions across the different combinations.

## DISCUSSION

It is generally expected that large populations are better able to adapt to an environmental challenge and reach higher fitness along more predictable evolutionary trajectories compared to small populations, because selection is more efficient and drift effects are dampened in large populations (Crow and Kimura, 1970). However, predictions from theory are ambiguous, especially for sexually reproducing species, and empirical data on the repeatability of evolutionary trajectories is focused mostly on asexual unicellular species. Using an obligately outcrossing line of the nematode *C. elegans*, we found similar average and maximum fitness across three different bottleneck treatments. A larger proportion of fitness variance could be attributed to drift in populations that had experienced a strong bottleneck, but neither the presence nor magnitude of the bottleneck significantly increased the variance in fitness.

Irrespective of bottleneck treatment, we observed that nearly all replicate *C. elegans* populations evolved higher relative fitness during experimental evolution under novel conditions. Given the relatively short time of our evolutionary experiment (∼21 generations) and the high genetic diversity of the *C. elegans* line used in this study (Noble et al., 2017), selection has most likely acted on standing genetic variation. Since a population bottleneck should reduce genetic variation, we had expected lower mean fitness and higher variance in fitness after evolution following a bottleneck compared to no bottleneck. These unexpected findings thus raise the question why fitness evolution in the nematodes was only marginally affected by population bottlenecks.

One potential explanation is that our bottleneck treatments did not reduce the effective population size and thus left genetic diversity unaffected. However, our fitness data support an effect of the bottleneck treatment pre-selection: variation in fitness between replicates as well as median fitness in the week 0 populations were lower after a bottleneck compared to populations that had no bottleneck treatment, although this was only statistically significant in the moderate bottleneck populations. Even though we cannot explain the observation of seemingly stronger effects on starting fitness in the moderate compared to the strong bottleneck result, it does not affect our conclusions from the experiments after 15 weeks. This is because (i) lower starting fitness would potentially lead to lower final fitness and higher fitness variance, neither of which we observed, and (ii) the only significant differences we observed at the end of the experiment were between the populations without bottleneck and the populations with strong bottleneck. In addition, when we partitioned the variance, we found an increased contribution of chance effects in the populations after a strong bottleneck, which is expected when genetic diversity is reduced. Jointly, these results support a biologically relevant effect of the bottleneck treatment on the amount of genetic variation available to selection.

An alternative explanation for our finding that all populations evolved higher relative fitness under novel conditions is that the adaptive potential of populations with reduced genetic variation is context dependent. For example, lab studies with *E. coli* populations have shown that bottleneck effects on the repeatability of fitness evolution depend on the traits that are under selection during adaptation, e.g. reduced repeatability across smaller *E. coli* populations under selection for antibiotic resistance (Windels et al., 2021) but not under selection for thermal tolerance (Wein and Dagan, 2019). Theoretical work has further shown that evolutionary predictability may not be uniformly influenced by effective population size, but that predictability is constrained in both very small and very large populations (Szendro et al., 2013). Lastly, sexual reproduction may modulate the effects of reduced genetic diversity on evolutionary repeatability in small populations (Weissman et al., 2010). Since we conducted our experiments with obligate outcrossing, sexual *C. elegans* populations, it is possible that the effects of population bottlenecks on fitness evolution were mitigated by recombination.

The genetic architecture may be a critical factor in predicting fitness evolution in relation to population bottlenecks. For example, in *C. elegans* epistatic interactions between genes underlying behavioral and fitness traits are wide-spread (Gaertner et al., 2012; Noble et al., 2017). Since epistasis is likely to reduce the effect of selfing on inbreeding depression (Abu Awad and Roze, 2020), these epistatic interactions may have evolved as a result of adaptation to a self-fertilizing life history with frequent cycles of exponential population growth followed by population crashes (Frézal and Félix, 2015). In addition, epigenetic changes may also contribute to the evolutionary responses observed here (Cavalli and Heard, 2019). Our common garden experimental design accounts for possible plastic and parentally (single-generation) heritable epigenetic changes, but the design does not account for any epigenetic changes that are stably inherited across multiple generations (Cavalli and Heard, 2019; Chey and Jose, 2022) and we can thus not exclude their role in driving fitness evolution. Both epistasis and epigenetic inheritance could buffer fitness evolution against reduced genetic diversity in small populations. We therefore suggest that our results fit a paradigm in which adaptive potential (and thus repeatability across replicated evolutionary events) does not unequivocally depend on effective population size, but that this relationship is shaped by the balance between diversity at neutral versus selected loci, by the strength of selection, and by the genetic architecture of selection responses (Bock et al., 2015; Carlson et al., 2014; Schrieber and Lachmuth, 2017).

In conclusion, we found that a strong population bottleneck in an obligate outcrossing line of *C. elegans* resulted in a higher contribution from drift and lower contribution from selection to fitness variation compared to populations that did not undergo a bottleneck. Importantly, due to our experimental setup we can exclude that the increased contribution from drift is due to (collinear) differences in census population size. The effects of bottlenecking on the evolution of fitness are marginal, as we observed only minor differences in fitness increase between treatments over the selection period, as well as in the repeatability of this fitness increase. Our results suggest a context-dependent relationship between genetic diversity, the effect of selection, and the predictability of evolutionary change.

## ACKNOWLEDGEMENTS

We would like to thank Juliane Teapal and the Utrecht University Large-Particle Flow Cytometry Facility (UU-LPC) for their help with the BioSorter, Suzanne Wiezer from Aquatic Ecology at NIOO-KNAW for using the Petri plate filling machine, and Sanne van der Steen. We also acknowledge Arjan de Visser, The Predicting Evolution consortium (Dries Bonte, Mirte Bosse, Steven Declerck, Marjon de Vos, Rampal S. Etienne, Steven Goossens, Martien Groenen, Paulien Hogeweg, Jan Kammenga, Ken Kraaijeveld, Martine Maan, Frederik Mortier, Ido R. Pen, Joost Riksen, Isabel Smallegange, Maurijn van der Zee, Sander van Doorn, Koen Verhoeven, Bregje Wertheim, Suzanne Wiezer, Lars E. Zandbergen), and the Origins Center for helpful discussions. This work was funded by The Dutch Research Council National Science Agenda NWA-ORC 400.17.606/4175 and a Flemish Research Foundation fellowship awarded to KB FWO-12T5622N.

## AUTHOR CONTRIBUTIONS

All authors conceived and designed the study; JM performed the experiments; KB and TB analyzed the data; KB and TB drafted the initial version of the manuscript and all authors contributed to later versions of the manuscript.

## DATA ACCESSIBILITY

The data and R code will be made available on Figshare upon acceptance.

