## Supplementary Information for "Population bottleneck has only marginal effect on fitness evolution and its repeatability in dioecious *C. elegans*"

Running title: Effect of genetic bottleneck on fitness evolution

### SUPPLEMENTARY METHODS

#### Study species and creation of the bottlenecked populations

The bacterivorous soil nematode *Caenorhabditis elegans* is highly suitable for experimental evolution given its small body size (adult females are approximately 1 mm), high fecundity (about 300 eggs per adult female per week), short generation time (can be as short as 50h), and easiness to maintain in the lab (Gray and Cutter, 2014; Stiernagle, 2006). Besides, frozen records can be created given that the first two larval stages can undergo cryopreservation (Gray and Cutter, 2014). The *C. elegans* D00 population from the Teotónio lab (IBENS, Paris) served as the initial source population for our experiment, which is a multiparent intercrossed population that is obligatorily outcrossing (Noble et al., 2017; Theologidis et al., 2014). Sex ratio in a dioecious *Caenorhabditis* species is expected to be 1:1 (Gray and Cutter, 2014) and fertility is not different from self-fertilized hermaphrodites (Theologidis et al., 2014).

The D00 source population was expanded on *E. coli* OP50 at 20°C and divided in aliquots. *E. coli* OP50 was maintained at 4°C. Bacterial lawns were added on the Nematode Growth Medium (NGM) (Stiernagle, 2006) plates by pipetting 50 µL of bacteria in L broth (10 g Bacto-tryptone, 5 g Bacto-yeast, 5 g NaCl, $H_2O$ to 1 liter; pH 7.0). To create the bottlenecked populations, five aliquots of the starting population were defrosted by holding the cryo-tubes in the hand. When thawed, the liquid was directly pipetted onto NGM plates to prevent oxygen depletion. This initial step was performed at 20°C with *E. coli* OP50 as a food source to avoid selection during the expansion. After six days, each aliquot was used to transfer 5 fertilized female nematodes (“strong bottleneck”) and 50 fertilized female nematodes (“moderate bottleneck”) by hand to two separate plates (see Fig. S1). These populations could grow for six days on NGM *E. coli* at 20°C before collecting the nematodes in Eppendorf tubes. Simultaneously with this expansion, the other five aliquots from the ancestral population were thawed and pipetted onto NGM *E. coli* at 20°C for six days for the “no bottleneck” treatment (Fig. S1). After growing, nematodes were extracted from the plates by rinsing the plates with S buffer (Stiernagle, 2006) and the nematodes were collected in 10 mL tubes. The density of the populations was measured for the fifteen populations to define the necessary volume to initiate the evolutionary experiment with 500 nematodes per plate. So, no bottleneck and bottleneck treatments always started with 500 nematodes in an expected 1:1 sex ratio, but for the no bottleneck treatment, these 500 nematodes were offspring of a diverse ancestral populations, while for the bottleneck treatments, these 500 nematodes were offspring of 50 or only 5 founder females. The females were chosen randomly from all available females that were adult but did not carry any embryo. We only selected females to avoid variation among bottleneck replicates by stochastically sampling different numbers of males and females: for example, offspring from1 male and 4 fertilized females will have a drastically higher effective size than offspring from 1 female and 4 males. Each treatment (no bottleneck, moderate bottleneck, and strong bottleneck) had five replicates consisting of three plates that were each initiated with 500 nematodes (Fig. S1). All populations of *C. elegans* were maintained on plates (⌀ 9 cm) with ±12 mL NGM. These NGM plates were poured using a Petri dish filling machine at NIOO-KNAW.

Novel conditions for experimental evolution

During the experimental evolution the *C. elegans* populations were grown on *Bacillus megaterium* (DSM No. 509), which is a gram-positive bacterium that is known to be hard-to-eat; *C. elegans* grows slower and avoids these bacteria when given a choice (Shtonda and Avery, 2006). *B. megaterium* was maintained at 4°C and bacterial lawns were added by pipetting 50 µL of bacteria in L broth on fresh NGM plates, which were then incubated at room temperature for 24 hours prior to transferring the worms.

For the experiment, the growing temperature was lowered to 16°C, which was done for experimental feasibility. The temperature reduction to 16°C may affect many processes such as metabolic functions and defense pathways (Gómez-Orte et al., 2018) and therefore constitute an additional selection pressure. Unexpectedly, 16S amplicon sequencing data from empty NGM plates revealed contamination of the plates (mainly bacteria from the genera *Serratia* and *Pseudomonas*). Although empty plates did not reveal any visual bacterial growth at room temperature and the same plates were used for all the treatments, this contamination may have induced an unanticipated additional selection pressure. Therefore, the effects of the three perturbations (novel food source, novel temperature, and plate contaminants) cannot be disentangled and are together considered as the novel conditions.

<u>Experimental set-up</u>

The experiment was initiated on fresh NGM plates with a lawn of *B. megaterium* and 500 nematodes per plate per replicate. For each replicate there were three plates that were mixed prior to the weekly transfer to sustain more genetic variation, larger population sizes, and to avoid the loss of a replicate if one plate would fail. Every week 500 nematodes were transferred by washing the plates, mixing the three plates per replicate, estimating the density of nematodes and pipetting the necessary volume to the new plates. At the beginning (week

0) and end (week 15) of the experiment, the nematodes were cryopreserved in a 12.5% glycerol liquid freezing solution using a MrFrosty (Thermo Fisher Scientific, Waltham, MA, USA) to gradually cool them to -80°C until they were needed for fitness assessments.

Fitness assessment

As a fitness proxy for each replicate nematode population, we used the population size after one week of growth on *B. megaterium* at 16°C (Fig. S2). One week of growth is identical to a single rearing step in the evolutionary experiment as explained in the experimental set-up. An increase in the numbers of nematodes at the end of a single rearing step is thus an estimate for an increase in fitness. We assume that a higher total number of nematodes after one week of growth for the final compared to the starting population is indicative of adaptation to the novel conditions.

The set-up of the fitness assessment is presented in Fig. S2. After thawing the sample, an expansion step is performed to increase the number of nematodes. For this, 250 μL of the thawed sample is pipetted on *E. coli* at 20°C to expand the population. The expansion was for one week to create a common garden and obtain sufficient nematodes for the start of the fitness assessment. The expansion may potentially select for nematodes that (still) feed efficiently on *E. coli*, but since *E. coli* are a very accessible prey to *C. elegans* across many different wild isolates, we do not expect this to be a significant selection pressure. After expansion, the population density is estimated (by averaging the number of nematodes in three droplets of 1 μL) to calculate the necessary volume to initiate the fitness assessment with 500 nematodes. Three technical replicates are made per sample. The population size of each fitness assessment is counted by extrapolating the density estimation (number of nematodes in 5 μL). We compared our counts with the counts from a flow cytometer

(BioSorter at Utrecht University) which resulted in a strong linear correlation (Fig. S3). For some populations, the fitness assessments were performed multiple times, these counts on the different measurement assessment days were used to quantify measurement error (Fig. S2).

FIGURES

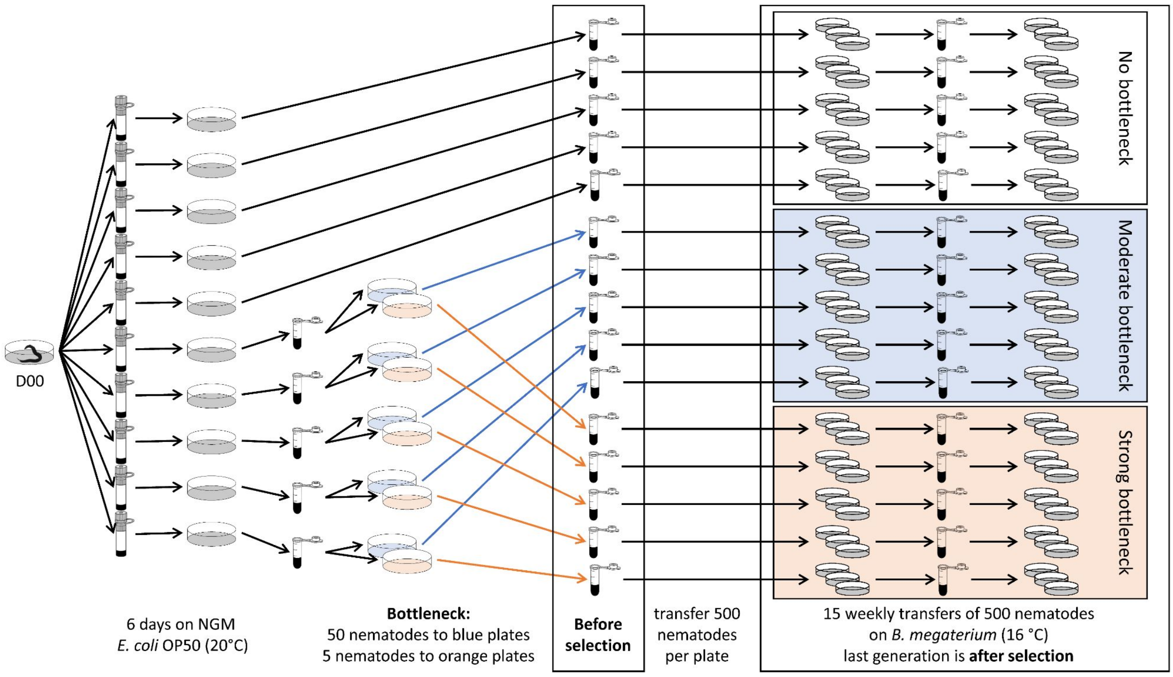


**Figure S1: General overview figure.** All treatments started from the same ancestral population that was expanded on ten petri dishes with NGM *E. coli* at 20°C. Bottlenecked populations were created from these expanded populations by transferring fifty nematodes (in blue; moderate bottleneck) or five nematodes (in orange; strong bottleneck). These populations ('before selection') were the start populations for the evolutionary experiment with fifteen weekly transfers onto fresh NGM *B. megaterium* plates at 16°C. Every time 500 nematodes were transferred per petri dish (with three petri dishes per replicate). The last generation is referred to as 'after selection'. There were five replicates per treatment.

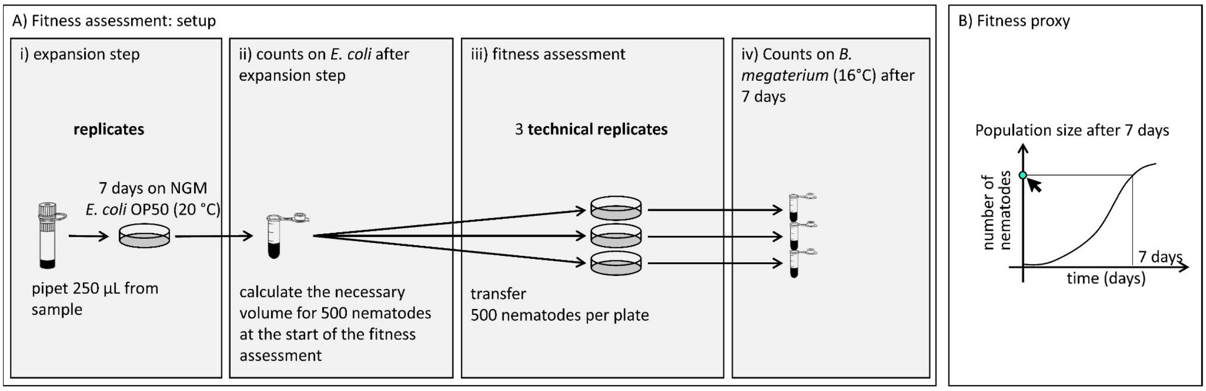


**Figure S2: Fitness assessment: setup and proxy.** A) Each replicate (five replicates before selection and five replicates after selection per treatment) undergoes an expansion step of one week to obtain sufficient nematodes for the fitness assessment (i). Based on the counts of the number of nematodes after the expansion step (ii) the required volume is quantified to initiate the assessment (iii), which is done by counting the number of nematodes after 7 days (iv) with three technical. B) The used fitness proxy is the nematode population size after seven days.

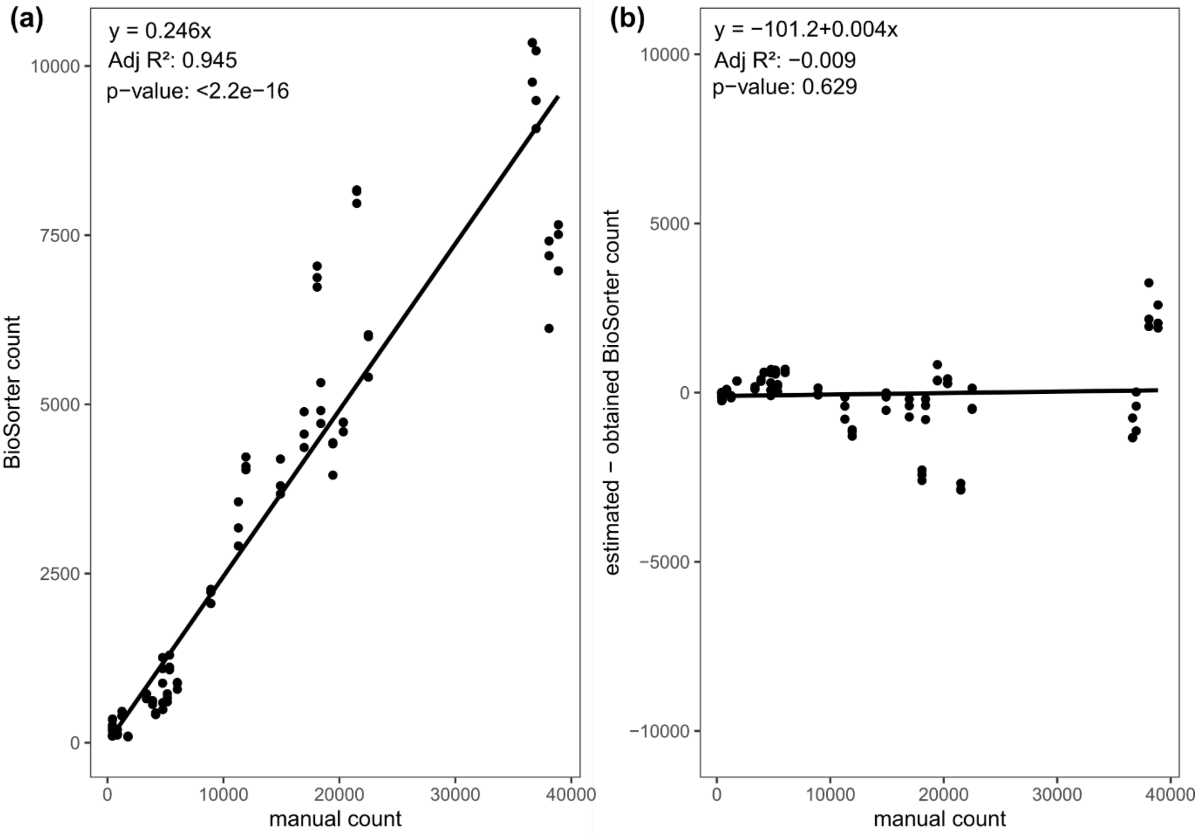

**Figure S3. Correlation between manual and BioSorter counts.** (a) The x-axis shows the manual counts, while the y-axis gives the results from the flow cytometer or BioSorter. Each dot represents the same technical replicate (see Fig. S2) for which the population size was assessed. The linear regression through the origin is indicated with the solid line and the equation is presented in the upper left corner. (b) The manual counts are given on the x-axis, while the y-axis represents the difference between the estimated counts based on the equation in (a) and the obtained counts from the flow cytometer

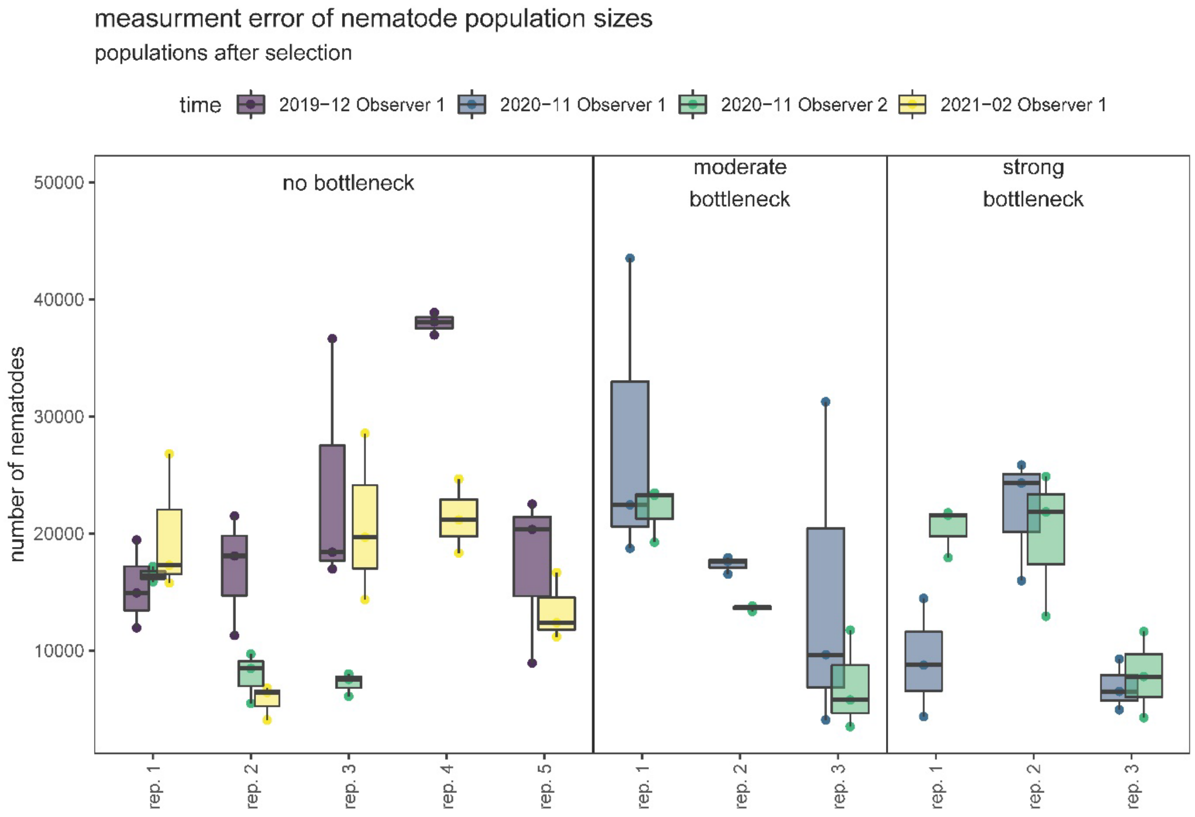


**Figure S4. Population size after 7 days of growth on *B. megaterium* before and after selection, partitioned across measurement days.** Each box-plot shows the distribution of three measurements taken at a given day (colors) by either observer 1 (purple, blue, yellow) or observer 2 (green, only for populations AFTER selection, for three (out of five) replicates per treatment (no, moderate, strong bottleneck). Labels: BB.C = backbone before selection,

BB.S = backbone after selection, PB.M = moderate population bottleneck (after selection), PB.S = strong population bottleneck (after selection). The figure again shows (just like in Fig. 1) that average but not maximum fitness across replicates is lower in the bottleneck treatment. Additionally, it provides a different visualization for the repeatability. It shows there is more relative variation due to chance (= differences between replicates within treatments after selection) in bottlenecked replicates compared to standard replicates. In the absence of a bottleneck, it seems the relative variation that can be attributed to selection (i.e. difference between before and after selection) is higher, but this will depend on what the measurements for week 0 for the bottleneck populations look like. Also, it seems that relative variation due to measurement error (i.e. differences between measurements taken on a given day) in bottlenecked replicates compared to standard replicates. These results will need to be confirmed analytically, using GLMs or sums-of-squares variance partitioning.

TABLES

**Table S1: Number of fitness assessment days and total measurements per treatment replicate.** The different columns indicate the treatments, weeks when the fitness was assessed during the experiment, the replicate numbers, fitness assessment days and total number of measurements.

| treatment | week | replicate | Number of fitness assessment days | measurements |
|---|---|---|---|---|
| no bottleneck | week 0 | **1** | 1 | 3 |
| | | **2** | 1 | 3 |
| | | **3** | 1 | 3 |
| | | **4** | 1 | 3 |
| | | **5** | 1 | 3 |
| | week 15 | **1** | 3 | 9 |
| | | **2** | 3 | 9 |
| | | **3** | 3 | 9 |
| | | **4** | 2 | 9 |
| | | **5** | 2 | 9 |
| moderate bottleneck | week 0 | **1** | 1 | 3 |
| | | **2** | 1 | 3 |
| | | **3** | 1 | 3 |

| | | | | |
|---|---|---|---|---|
| | | **4** | 1 | 3 |
| | | **5** | 1 | 3 |
| | **week 15** | **1** | 2 | 6 |
| | | **2** | 2 | 6 |
| | | **3** | 2 | 6 |
| | | **4** | 1 | 3 |
| | | **5** | 1 | 3 |
| **strong bottleneck** | **week 0** | **1** | 1 | 3 |
| | | **2** | 1 | 3 |
| | | **3** | 1 | 3 |
| | | **4** | 1 | 3 |
| | | **5** | 1 | 3 |
| | **week 15** | **1** | 2 | 6 |
| | | **2** | 2 | 6 |
| | | **3** | 2 | 6 |
| | | **4** | 1 | 3 |
| | | **5** | 1 | 3 |
